# Developmental dynamics are a proxy for selective pressures on alternatively polyadenylated isoforms

**DOI:** 10.1101/737551

**Authors:** Michal Levin, Harel Zalts, Natalia Mostov, Tamar Hashimshony, Itai Yanai

## Abstract

Alternative polyadenylation (APA) leads to multiple transcripts from the same gene, yet their distinct functional attributes remain largely unknown. Here, we introduce APA-seq to detect the expression levels of APA isoforms from 3’-end RNA-Seq data by exploiting both paired-end reads for gene isoform identification and quantification. Applying APA-seq, we detected the expression levels of APA isoforms from RNA-Seq data of single *C. elegans* embryos, and studied the patterns of 3’ UTR isoform expression throughout embryogenesis. We found that global changes in APA usage demarcate developmental stages, suggesting a requirement for distinct 3’ UTR isoforms throughout embryogenesis. We distinguished two classes of genes, depending upon the correlation between the temporal profiles of their isoforms: those with highly correlated isoforms (HCI) and those with lowly correlated isoforms (LCI) across time. This led us to hypothesize that variants produced with similar expression profiles may be the product of biological noise, while the LCI variants may be under tighter selection and consequently their distinct 3’ UTR isoforms are more likely to have functional consequences. Supporting this notion, we found that LCI genes have significantly more miRNA binding sites, more correlated expression profiles with those of their targeting miRNAs and a relative lack of correspondence between their transcription and protein abundances. Collectively, our results suggest that a lack of coherence among the regulation of 3’ UTR isoforms is a proxy for selective pressures acting upon APA usage and consequently for their functional relevance.

## Introduction

Alternative polyadenylation (APA) is a crucial regulatory mechanism – widespread and conserved across all eukaryotes – that diversifies post-transcriptional regulation by selective mRNA-miRNA and mRNA-protein interactions (Ozsolak et al. 2010; Jan et al. 2011; Derti et al. 2012; Smibert et al. 2012; Ulitsky et al. 2012; Velten et al. 2015; Hu et al. 2017). APA plays an important role in a vast variety of biological processes, such as maternal to zygotic transition, cell differentiation, and tissue specification (Wormington 1994; Tadros et al. 2007b; Tadros and Lipshitz 2009) and exerts a tremendous influence over gene expression, the transcript’s cellular localization, stability and translation rate (Mazumder et al. 2003; Keene 2007; Lutz and Moreira 2011; Elkon et al. 2013; Tian and Manley 2017). Since first discovered in the immunoglobulin M and the DHFR genes (Alt et al. 1980; Early et al. 1980; Rogers et al. 1980; Setzer et al. 1980), technological advances and high-throughput sequencing techniques have made it clear that APA is more widespread than initially thought; 30-70% of the genes undergo APA in diverse species (Mangone et al. 2010; Ozsolak et al. 2010; Jan et al. 2011; Derti et al. 2012; Smibert et al. 2012; Ulitsky et al. 2012; Velten et al. 2015). During the last decade, widespread APA alterations were detected across different tissues and across distinct stages of embryogenesis (Tian et al. 2005; Tadros et al. 2007a; Wang et al. 2008; Ji et al. 2009; Li et al. 2012; Smibert et al. 2012; Ulitsky et al. 2012; Blazie et al. 2015; Blazie et al. 2017; Hu et al. 2017; Khraiwesh and Salehi-Ashtiani 2017). However, beyond these classifications and functional characterization of a short list of single gene APA alterations (Chen et al. 2017) the global functional significance of APA remains largely elusive. In light of the vast amount of alternative 3’UTR isoforms that have been detected during the past few years it would be beneficial to be able to enrich for those alterations which have functional consequences.

*C. elegans* is a convenient model organism for studying APA and gene expression during embryogenesis, since the cell lineage is invariant and has been fully traced (Sulston and Horvitz 1977; Kimble and Hirsh 1979). *C. elegans* was the first multicellular organism to have a fully sequenced genome (Consortium 1998); and its transcriptome has also been well characterized (McKay et al. 2003; Shin et al. 2008; Hillier et al. 2009; Ramani et al. 2009; Lamm et al. 2011; Grün et al. 2014). During embryonic development, the *C. elegans* transcriptome is highly dynamic; in early stages it is comprised mostly of maternal transcripts, but as development proceeds, zygotic transcription commences, and maternally supplied transcripts undergo degradation (Newman-Smith and Rothman 1998; Tadros et al. 2007b; Tadros and Lipshitz 2009; Walser and Lipshitz 2011). Our previous results detected many genes with a dynamic overall gene expression profile throughout embryogenesis, as well as genes with constitutive levels of expression (Levin et al. 2012).

CEL-Seq is a sensitive multiplexed single-cell RNA-Seq method (Hashimshony et al. 2012; Hashimshony et al. 2016). One important feature of the CEL-Seq method is that it is restricted to studying the 3’end of the transcriptome and thus measures overall expression levels; typically collapsing the various isoforms produced by a gene to one summary profile. While this has been a useful simplifying criterion, it does ignore possible dynamic profiles across different splicing isoforms of a particular gene. An important advantage of the CEL-Seq 3’end bias though is that this information can be used to detect and quantify alternative polyadenylation patterns *i.e.* 3’ UTR isoforms of the same gene, as we propose here.

Here, we studied APA profiles in individual *C. elegans* throughout embryogenesis using APA-seq, an approach to detect alternative polyadenylation profiles at a genomic scale. APA-seq is based on CEL-Seq but further exploits the information from both paired-end reads allows for gene identification from Read 2 and the exact location of polyadenylation from Read 1. Combining this information empowers quantitative expression level assessment globally at both the gene and APA isoform level. Using this approach, we delineated two groups of genes, those with highly and lowly correlated 3’ UTR isoform groups (HCI and LCI), respectively. We detected unique regulatory features between these groups which supports the notion that variants across these two groups are under distinct selective biases. Genes with uncorrelated 3’ UTR isoform expression (LCI) are predicted to have the highest miRNA regulation compared to genes with well-correlated 3’ UTR isoforms. Integrating extensive previously published embryonic mRNA and protein expression datasets (Mangone et al. 2008; Grün et al. 2014), we also found the lowest correlation between total mRNA transcript and protein levels in genes with dynamic 3’ UTR isoform expression (LCI). Extending this analysis to *Drosophila melanogaster* and *Xenopus laevis* datasets we found a consistent relationship (Graveley et al. 2011; Casas-Vila et al. 2017; Sanfilippo et al. 2017; Zhou et al. 2019; Peshkin et al. 2015). Together, our results suggest that genes of the HCI and LCI groups experience distinct regulatory pressures upon their alternatively polyadenylated isoforms.

## Results

### APA-seq identifies expression levels of alternative polyadenylation isoforms

Paired-end reads generated by CEL-Seq and CEL-Seq2 contain the sample-specific barcode on Read 1 and the sequence identifying the transcript on Read 2 (Hashimshony et al. 2012; Hashimshony et al. 2016) (Fig. 1A). Typically, in CEL-Seq only Read 2 is used for measuring gene expression levels. However, since CEL-Seq sequences the 3’ ends, it can be used in principle to identify 3’ UTR isoforms. Using Read 2 for this purpose is challenging due to the uneven sizes of the inserts in the sequencing library, producing a smear of mapped reads, which makes distinguishing different 3’ UTR isoforms of the same transcript impossible in many cases (Fig. 1B, red peak). However, we noted that Read 1 includes the actual 3’end of the transcript (Fig. 1A), located just upstream of the polyadenylation site (Fig. 1B, black peak). In CEL-Seq, this region follows 24 Ts used for capturing the polyA tail of the transcript, and the sequencing quality is relatively poor after this low complexity region rendering conventional mapping impossible (Supplemental Fig. S1A,B). To overcome this, we found that the sequence is still of sufficient quality when mapping is performed using relaxed parameters and restricted to a particular region of the genome. In summary, while permissive Read 1 mapping matches many genomic loci due to its poor quality, it maps uniquely when restricted to the sequence of a particular gene, whose identity is detected by using Read 2.

**Fig. 1.**
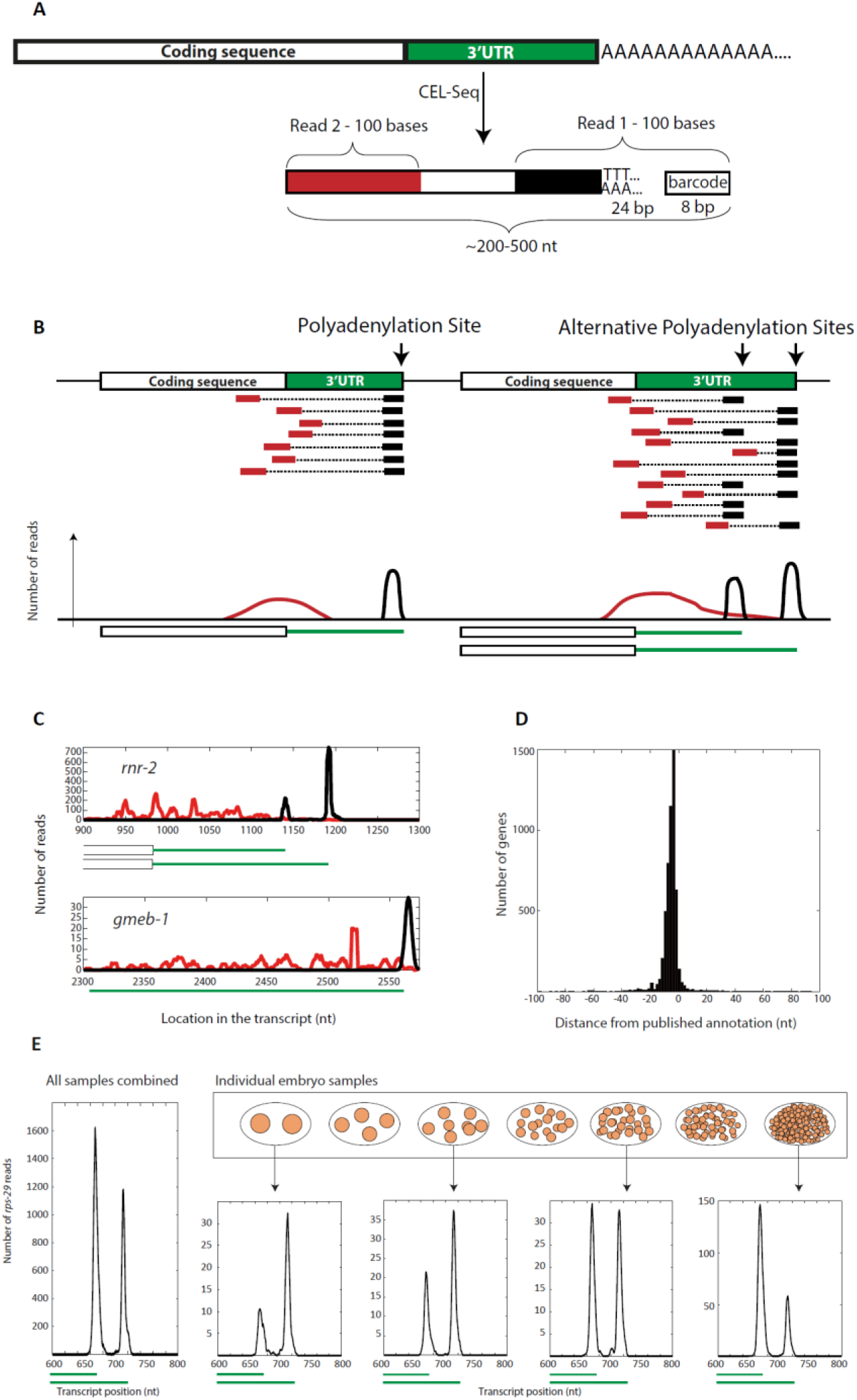
APA-seq measures expression levels of distinct 3’ UTR isoforms in individual *C. elegans* embryos. (A) APA-seq identifies gene expression levels for distinct alternatively polyadenylated isoforms. APA-seq is an adaptation of the CEL-Seq method which utilizes paired-end reads: Read 1 contains a sample-specific barcode while Read 2 identifies the transcript. If sequenced long enough (100bp in our case), Read 1 also provides information on the exact location of the polyadenylation site. APA-seq thus uses Read 2 to identify the expressed gene and then maps Read 1 to the gene specific region, thus enabling unique mapping in spite of the low sequencing quality that results from sequencing through the low-complexity poly-T region. (B) Mapping Read 2 sequences results in a wide distribution within the gene (red peak) due to the fragmentation step of library preparation. However, Read 1 sequences all map to the site immediately upstream of the poly-A tail thus producing a clear peak (black peak) when mapped to the gene sequence and revealing exact polyadenylation sites. The white boxes at the bottom of the distribution plots mark the coding sequence, while the green lines indicate the determined 3’ UTR regions. (C) Using the APA-seq method enables detection of the exact location of polyadenylation sites in *C. elegans*. The green lines at the bottom of each plot mark previously annotated 3’ UTRs (Mangone et al. 2008) for the indicated gene, showing good agreement with the APA-seq Read 1 peaks (black). (D) Global comparison of the detected polyadenylation sites using APA-seq to the *C. elegans* UTRome annotation (Mangone et al. 2008) shows high consistency between the two datasets. (E) The expression of two unique 3’ UTR isoforms for the *C. elegans rps-29* gene throughout embryogenesis. Although in total (leftmost panel) the 3’ UTR variants show equal expression, examining expression across developmental stages shows predominant expression of the shorter 3’ UTR variant during early embryogenesis, and the inverse later in development.

We thus devised the APA-seq approach to study 3’ UTR isoforms using both Read 1 and Read 2 information and applied it to study the expression of alternative polyadenylated isoforms during early embryogenesis in the nematode *C. elegans*. For this, we used a dataset previously published by our lab in which embryos were individually collected and sequenced throughout embryogenesis (Levin et al. 2016). Pooling together all samples, we detected distinct 3’ UTR isoforms for 5,336 genes using APA-seq, after removing possible artifacts caused by internal priming (Gruber et al. 2016) (see Methods). For example, the *C. elegans* genes *gmeb-1* and *rnr-1* (Fig. 1C) show a smear of Read 2 mappings (shown in red), while Read 1 mappings form distinct peaks (shown in black) identifying previously characterized polyadenylation sites (shown in green) (Mangone et al. 2008) (see also Supplemental Fig. S1C).

Overall, we detected multiple 3’ UTR isoforms for 14% of the expressed genes in our dataset (746 out of 5336 genes). Of these, less than 1% have more than two isoforms (Supplemental Fig. S1D). This set is not comprehensive to all possible 3’ UTR isoforms produced by the *C. elegans* genome given that we only examined 430 minutes during embryogenesis, and the stringent thresholds set in our bioinformatics pipeline (in terms of mapping parameters, mapping level filtering and spurious site removal, see Methods). We assayed the accuracy of the isoforms detected by comparing our polyadenylation sites with a known repository of 3’ UTR annotations in *C. elegans* (Mangone et al. 2008) and found highly concordant profiles, with 95% of sites corresponding to well-established annotated sites (Fig. 1D, Supplemental Table S1). The remaining 5% show significantly lower expression levels than the annotation overlapping 3’ UTR isoforms (Supplemental Fig. S1E) and we therefore excluded these from further analyses by expression level filtering. To study the temporal dynamics of 3’ UTR isoform expression, we classified the reads according to their embryo of origin (Fig. 1E, Supplemental Table S1). As an example, two isoforms were identified for *rps-29*, which encodes a ribosomal protein subunit (Kamath et al. 2003). Interestingly, while the sum of expression of both *rps-29* isoforms is roughly constant over time, the long 3’ UTR isoform is predominantly expressed in the early stages while the shorter is expressed in later embryos (Fig. 1E). Correlating the expression levels of all 3’ UTR isoforms across stages, we found that successive stages (near-replicates) show high correlations thus highlighting the reproducibility of the data (Supplemental Fig. S2). We conclude that while CEL-Seq is typically used to assay overall expression levels with the mapping location typically ignored, processing the reads using APA-seq can identify the exact locations of alternative polyadenylation sites and the expression levels of distinct 3’ UTR isoforms.

### Alternative polyadenylation profiles throughout C. elegans embryogenesis

Our dataset enabled us to study overall patterns of 3’ UTR isoform usage throughout early development in individual embryos. For each gene we first computed the ratio of expression between its 3’ UTR isoforms throughout the time-course. Interestingly, the clustering of stages according to these profiles revealed that adjacent developmental stages show concordant 3’ UTR isoform usage patterns which group into distinct periods with diverging APA dynamics (Fig. 2A). The four groups correspond to previously characterized developmental periods: maternal degradation, early period of extensive proliferation and specification, the mid-embryonic transition period and the subsequent period of morphogenesis (47). The observation that different periods have different corresponding patterns of 3’ UTR isoforms may reflect that each of these has a distinct functional requirement. Between adjacent periods we found that the direction and level of change of polyadenylation site usage generally shows a broad burst of shortening especially between the proliferative and mid-embryonic transition periods (Fig. 2B). This is consistent with previous work indicating that proliferative states show an overall trend for 3’ UTR shortening (Tian and Manley 2017). Overall, we identified that 89% of dynamic 3’ UTR isoform genes show APA switches in at least one of the period switches indicated (Fig. 2C).

**Fig. 2.**
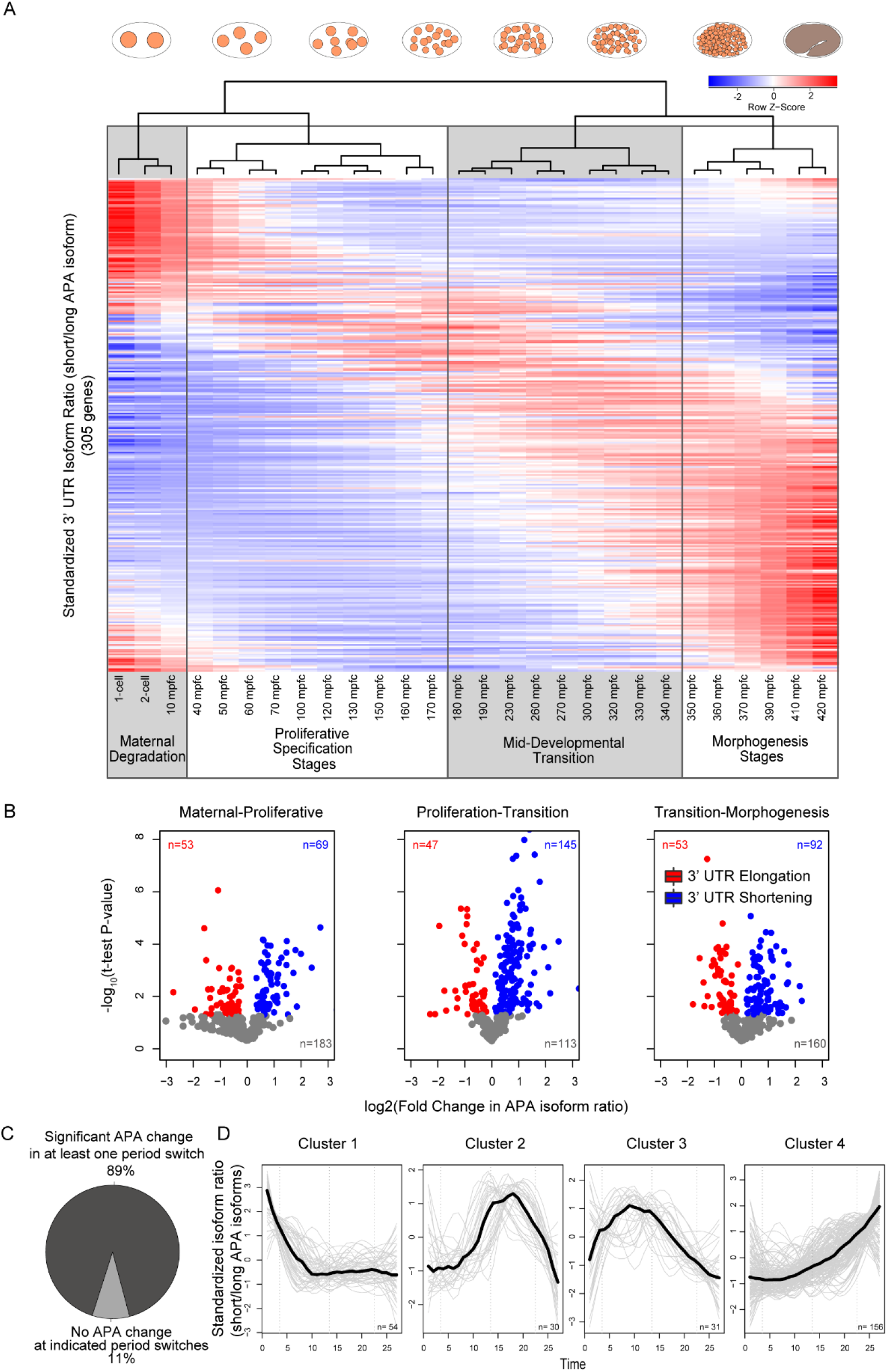
3’ UTR isoform expression throughout *C. elegans* embryogenesis. (A) Heatmap showing the relative expression of 3’ UTR isoforms for 305 genes which passed overall expression and 3’ UTR isoform dynamics threshold throughout *C. elegans* embryogenesis. Each row in the heatmap corresponds to a gene and indicates the ratio between the expression of its short and long isoforms. Red and blue indicate the maximum and minimum 3’ UTR ratio for each gene, respectively. White and grey shadowed boxes indicate sets of developmental stages with similar isoform usage (based on Ward clustering, see clustergram on top of the heatmap) and identify the periods of major isoform switches. mpfc = minutes past four-cell stage. (B) Studying the level of 3’ UTR isoform changes across development. The plots indicate the fold-change and *P*-value of difference in the isoform ratios for pairs of successive periods as defined by Ward clustering in A. Red and blue coloring indicates significant elongation and shortening events, respectively. Grey coloring indicates insignificant changes. (C) Most of significant 3’ UTR isoform switches occur between the identified developmental switch periods. 89% (271) of the genes show significant changes in 3’ UTR usage at least one of the identified period switches. (D) Clusters of significant 3’ UTR isoform changes indicate four main kinetics – almost constant elongation or shortening over time (Cluster 1 and 4, respectively) or peaking shortening during mid-developmental transition and proliferative periods (Cluster 3 and 4). The thick black line indicates the mean 3’ UTR isoform ratio profile of all genes in the cluster and individual genes profiles are shown as thin grey lines.

Clustering the 3’ UTR isoform ratio profiles throughout the time-course, we identified four main distinct clusters (Fig. 2D); genes whose 3’ UTRs elongate or shorten over time (Cluster 1 and 4, respectively) and genes whose 3’ UTRs shorten transiently during the proliferation period (Cluster 2 and 3). The majority of genes though show continuous shortening of their 3’ UTR regions with progressing development (cluster 4). Interestingly, these genes show enrichments for MAP kinase cascade, morphogenesis and neuronal differentiation (Supplemental Fig. S3). All these functions are crucial components of the switch from germ cell biology to proliferative and differentiation processes. In summary, we show that the 3’ UTR dynamics detected by APA-seq reflect characteristic functionalities of embryonic development and although the significance of single events is not clear yet the overall contribution of APA events to the general regulatory states involved in these events might be high or alternatively present a side effect of the vast changes occurring at the other levels of gene expression regulation.

### Genes with constitutive total expression are enriched for dynamic 3’ UTR isoform expression

Examining the temporal profiles, we found that 3’ UTR isoforms of a particular gene may exhibit striking dynamics throughout development, while the overall total expression for the gene may be uniform (Fig. 3A). To study this systematically, we defined the dynamic range of a gene’s overall expression profile as the fold differences between the maximum and minimum expression values throughout the time-course. We found a positive correlation between the dynamic overall expression range of a gene and the correlation among its isoforms (Fig. 3B; *r*=0.94, *P*=0.03, 2^nd^ degree polynomial regression test, N=746). Similar results were obtained using the interquartile range (IQR) of the time-course overall expression levels as a proxy for expression dynamicity (r=0.98, P=0.04, 3^rd^ degree polynomial regression test, N=746). This result provides evidence for the notion that apparently constitutive genes can be highly dynamic at the level of their individual 3’ UTR isoforms.

**Fig. 3.**
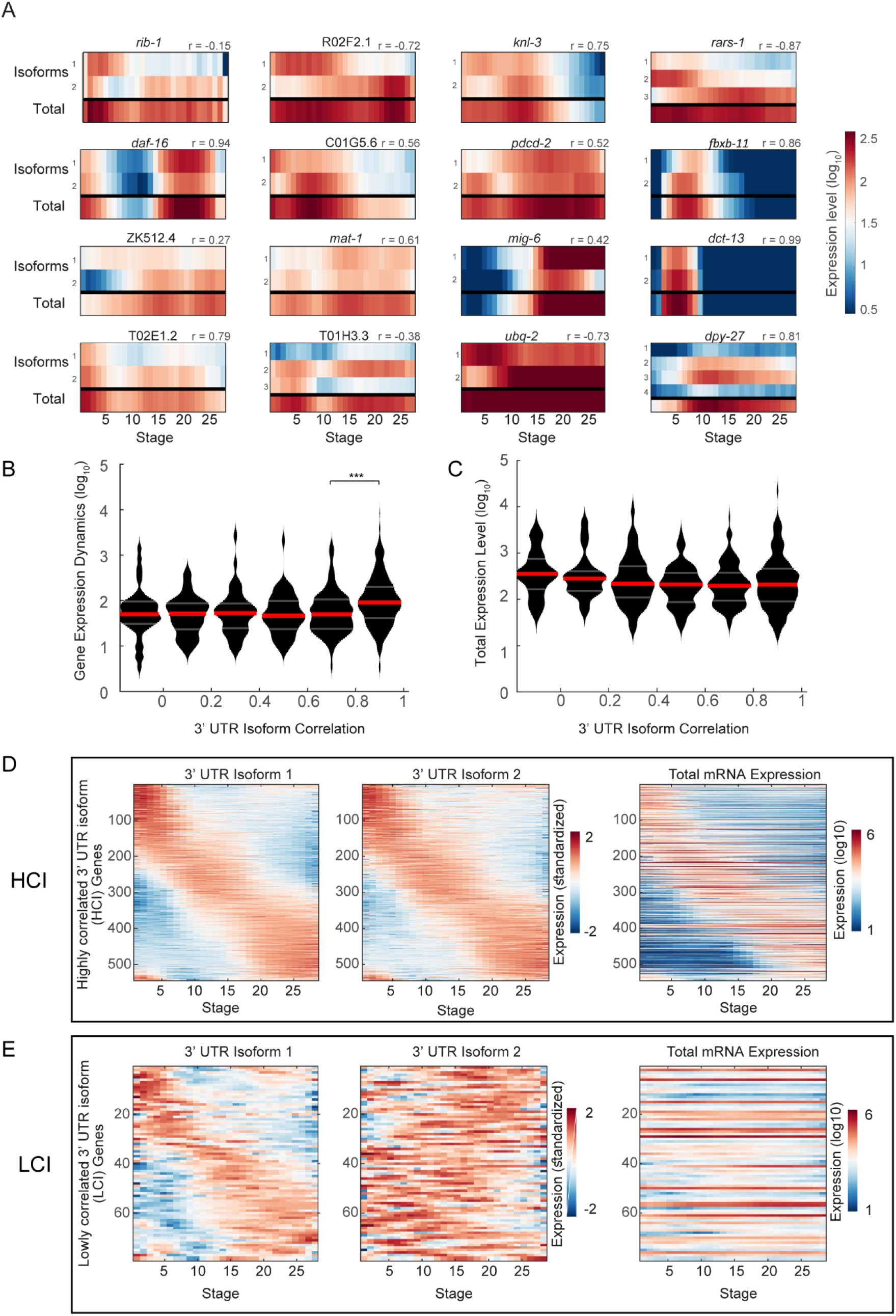
Genes with constitutive overall expression have dynamically expressed 3’ UTR isoforms. (A) Expression heatmaps indicating 3’ UTR isoform expression for 16 genes with multiple isoforms displaying varying levels of correlation between the expression of their 3’ UTR isoforms. The Pearson correlation coefficient *r* for each of the genes displayed is indicated at the top of each heatmap. (B) Relationship between 3’ UTR isoform expression correlations and the overall expression dynamics. Genes whose 3’ UTR isoform expression levels are correlated are more dynamic in their overall expression (*r*=0.94, *P*=0.03, 2^nd^ degree polynomial regression test). Dynamics for each gene is defined as the fold differences between its maximum and minimum expression values throughout the time-course. Red and grey horizontal bars represent the median and the interquartile ranges of the data, respectively. The last bin with highest 3’ UTR isoform correlations exhibit significantly higher overall expression dynamics than the preceding bin (P<10^−20^, Mann-Whitney test). (C) Relationship between 3’ UTR isoform expression correlations and the respective overall total mRNA expression levels. Genes whose 3’ UTR isoform expression levels are uncorrelated show significantly higher overall expression levels (*r*=0.99, *P*=0.002, 2^nd^ degree polynomial regression test). Red and grey horizontal bars represent the median and the interquartile ranges of the data, respectively (D) Heatmaps in the two leftmost panels show standardized expression of the 3’ UTR isoforms of 545 genes belonging to the group of genes whose 3’ UTR isoform’s expression show high correlation (HCI genes). The right panel shows a heatmap of the total mRNA expression (on a log10 scale) of the same genes confirming that genes with highly correlated isoform expression are also dynamically expressed. The time points are the same as indicated in Fig. 2A. (E) Same as D for 79 genes belonging to the group of genes whose 3’ UTR isoform’s expression show low correlation (LCI genes). Genes with distinct expression profiles of 3’ UTR isoforms appear constitutively expressed when examined at the total gene expression level.

Gene expression levels may explain this correlation, if genes with less dynamic behavior are also lowly expressed, and in turn may have noisier and uncorrelated 3’ UTR isoform profiles. To control for this possibility, we asked if there is a trend in expression levels according to the correlation among isoforms (Fig. 3C). Interestingly, genes with non-congruent APA behavior also show higher expression levels overall (Fig. 3C; *r*=0.99, *P*<0.002, 2^nd^ degree polynomial regression test, N=746) eliminating expression noise as a confounding factor.

To further examine this phenomenon, we visualized the dynamics of 3’ UTR isoform expression and total expression in genes with highly or lowly correlated isoforms. The 746 genes with multiple 3’ UTR isoforms displayed a variety of isoform expression correlations (Supplemental Fig. S4). More than 70% of the genes (545 genes) have highly correlated 3’ UTR isoform expression (*r*>0.7), while 11% of the profiles (79 genes) are lowly correlated (*r*<0.3). We henceforth refer to the two gene sets of highly and lowly correlated 3’ UTR isoforms, as HCI and LCI, respectively. As the heat maps in Fig. 3D-E show, we found dynamic expression for both the HCI and LCI groups at the 3’ UTR isoform level. As expected, the total gene expression for the correlated isoforms is dynamic, recovering the dynamics of the isoforms. Conversely, the total expression of the LCI genes mostly corresponds to profiles with constitutive overall expression. Such profiles are frequently attributed as housekeeping profiles, however as our analysis reveals, at the 3’ UTR isoform level they may be very dynamic. Thus, by de-convolving the total expression into profiles of distinct 3’ UTR isoforms we were able to extract a new layer of information from this dataset.

Delineating the LCI and HCI groups, led us to hypothesize that 3’ UTR isoforms from the former group are under stronger selective pressures, and are consequently more likely to be functionally different. We thus set out to test this hypothesis by studying the post-transcriptional and translational regulatory characteristics of these two gene sets.

### LCI genes contain more miRNA binding sites and correlate with miRNA expression

The 3’ UTR region is known to be a locus of considerable post-transcriptional regulation (Barrett et al. 2012; Pichon et al. 2012), and the role of miRNAs in this regulation is well evidenced (Ambros 2004; Bartel 2004). We thus searched for evidence that our detected 3’ UTR isoform expression profiles are regulated by miRNAs. Specifically, we asked if genes with a different number of 3’ UTR variants (single vs. multiple) and different 3’ UTR isoform expression correlations (HCI vs. LCI) have distinguishing sequence properties related to miRNA regulation (Fig. 4A). We first counted the number of basic miRNA seed matches in the 3’ UTR sequence of the different groups (Peterson et al. 2014). We found that genes with more than one 3’ UTR variant have significantly more miRNA binding sites than genes with a single 3’ UTR isoform (*P*<10^−12^, Mann-Whitney test, Fig. 4B). Genes with multiple variants also have significantly longer 3’ UTRs (*P*<10^−28^, Mann-Whitney test, Fig. 4C), though the number of miRNA binding sites per base is not different across the groups (Supplemental Fig. S5). Thus, genes with multiple variants are predicted to be regulated more actively by miRNAs.

**Fig. 4.**
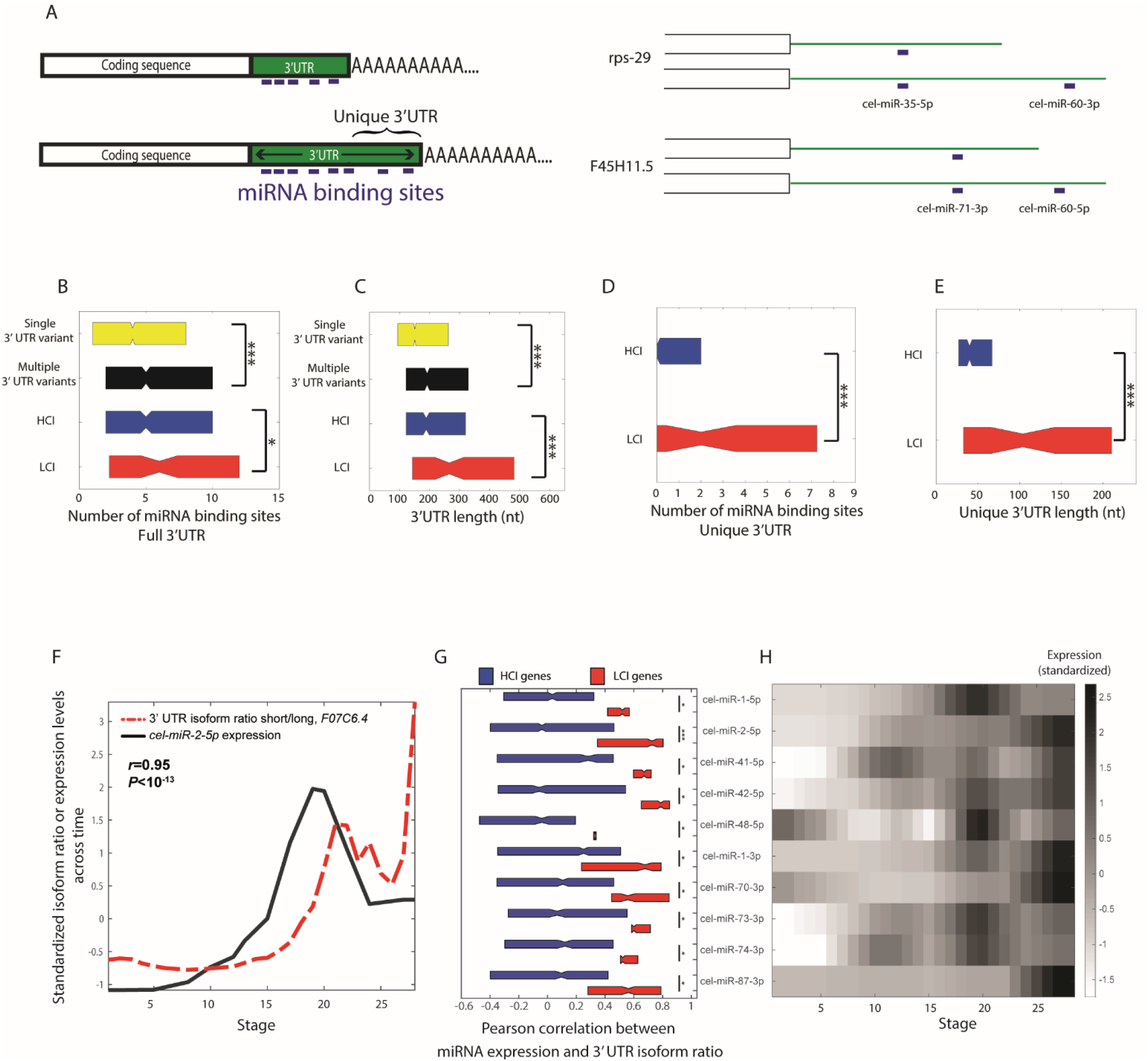
Genes with multiple, lowly correlated 3’ UTR variants show evidence for increased miRNA regulation. (A) The left panel shows a schematic representation of the miRNA analysis. The length of the 3’ UTR region, the number of basic miRNA seed matches, and the 3’ UTR region unique to the longer 3’ UTR isoform were considered. The right panel shows two examples of genes and their miRNA binding sites. (B) Boxplots indicating the number of miRNA binding sites in the full 3’ UTR regions of genes with single or multiple 3’ UTR isoforms, highly correlating (HCI) or lowly correlating isoforms (LCI). Genes with multiple 3’ UTR variants have significantly more miRNA targets than genes with a single variant (*P*<10^−12^, Mann-Whitney test). Between multiple 3’ UTR isoform genes, LCI genes have significantly more miRNA binding sites than HCI genes (*P*<0.02, Mann-Whitney test). (C) Same as B for the full 3’ UTR lengths of genes. Genes with multiple 3’ UTR variants have significantly longer 3’ UTRs than single isoform genes (*P*<10^−28^, Mann-Whitney test). Within multiple isoform genes, LCI genes have significantly longer 3’ UTRs than HCI genes (*P*<10^−3^, Mann-Whitney test). (D) Boxplots indicating the number of miRNA binding sites in the unique 3’ UTR region of the longer 3’ UTR isoform across HCI and LCI genes. LCI genes have significantly more miRNA binding sites in their unique 3’ UTR region than HCI genes (*P*<10^−3^, Mann-Whitney test). (E) Boxplots indicating the length of the unique 3’ UTR region across genes HCI and LCI genes. LCI genes have significantly longer unique 3’ UTR regions than HCI genes (*P*<10^−4^, Mann-Whitney test). (F) Correlating expression of miRNA expression with expression ratio dynamics between 3’ UTR isoforms. The black line shows the expression profile of *cel-miR-2-5p* miRNA throughout the developmental time-course. Depicted in red is the ratio between the expression profiles of the two 3’ UTR isoforms (short/long) of the *F07C6.4* gene. The Pearson correlation coefficients between 3’ UTR isoform ratio and miRNA expression of all target genes were used for the analysis shown in G and H. (G) Shown are the top ten miRNAs showing a significant difference between the HCI and LCI gene groups (out of 64 miRNAs with dynamic expression-statistics of all miRNAs can be found in Supplemental Table S2). All ten exhibit a positive median correlation between miRNA expression and the 3’ UTR isoform expression ratio of genes it is predicted to bind (as in F). Further, this correlation is significantly higher in LCI than in HCI genes. (H) Heatmap of the standardized expression dynamics of the ten indicated miRNAs which show differences between HCI and LCI genes across development.

Comparing the number of binding sites between the HCI and LCI groups, we found that the latter group has significantly more miRNA binding sites in their 3’ UTR (*P*<0.02, Mann-Whitney test, Fig. 4B). We further studied this group of genes by examining the sequence that is unique to the longer 3’ UTR isoform (Fig. 4A). Comparing between the HCI and LCI groups, we found that the latter have significantly more miRNA binding sites in their unique 3’ UTR region (*P*<10^−3^, Mann-Whitney test, Fig. 4D). This is not because LCI genes have denser distribution of miRNAs, rather their unique 3’UTR regions are significantly longer than in HCI genes (P<10^−4^, Mann-Whitney test, Fig. 4E), thus they tend to carry more potential miRNA binding sites. These results implicate a role for miRNAs in the differences we see between the genes sets of correlated and uncorrelated isoform genes.

To further examine the effect that miRNA regulation exerts on HCI and LCI genes, we computed correlations between the expression profiles of all dynamically expressed miRNAs (Avital et al. 2017) and the 3’ UTR isoform expression ratio of the genes they regulate. For example, *cel-miR-2* is conserved in *C. briggsae* as well as in *Drosophila*. Predicted targets of *mir-2* are enriched for genes involved in neural development (Marco et al. 2012). Expression of *miR-2* has been detected at all life stages, most abundantly in the L1 larval stage (Marco et al. 2012). Consistently, we detected expression in late embryogenesis in our miRNA expression data (Fig. 4F and 4H) and interestingly, its profile has a correlation of 0.94 (*P*<10^−13^) with the ratio of the 3’ UTR isoform expression of its target gene *F07C6.4*, a gene which is enriched in the germ line, germline precursor cell, the body wall musculature and in the PVD and OLL neurons (Smith et al. 2010; Lee et al. 2017). Figure 4G shows the distributions of correlation coefficients for dynamically expressed miRNAs with the 3’ UTR isoform expression ratio of their targets in the HCI and LCI groups. These ten miRNAs were selected based upon the most significant difference between the target correlations of the HCI and LCI gene groups (out of 64 miRNAs with dynamic expression-statistics of all miRNAs can be found in Supplemental Table S2). We found that in all ten the correlations are significantly higher in LCI than in HCI genes. For example, *cel-miR-2-5p* shows significantly higher correlations with the expression ratio of the isoforms of all the genes it potentially binds and regulates in the LCI genes (*P*<10^−3^, Mann-Whitney test, Fig. 4G). This analysis suggests that miRNAs play a major role in regulating genes with lowly correlated 3’ UTR isoforms (LCI) during *C. elegans* embryogenesis.

### LCI genes exhibit lower mRNA-protein correspondences

Beyond regulation by miRNA, control of translation efficiency constitutes another level of post-transcriptional regulation. 3’ UTR regions are known preferential targets for RNA binding proteins that regulate translation in terms of localization and efficiency (Szostak and Gebauer 2013; Zhao et al. 2014; Berkovits and Mayr 2015). Indeed, mRNA and protein levels are notoriously lowly correlated (Grün et al. 2014). We reasoned that one possible explanation for the low correspondence of some genes may follow from the fact that different 3’ UTR isoforms may exhibit different translation efficiencies. Thus, we predicted that LCI genes would have a worse correspondence – relative to HCI genes – between transcription and protein levels. To test this, we turned to a previously published mRNA and protein *C. elegans* time-course (Grün et al. 2014) and examined the distribution of correlations between mRNA and protein abundances across the HCI and LCI groups. We detected that LCI genes show lower correlations between mRNA and protein abundances, relative to the HCI genes (Fig. 5A; *P*=0.07, Mann-Whitney test; N=188, 30 for HCI and LCI, respectively). This limited significance may be due the low number of detected LCI genes following the shortness of the time-course, and the restricted protein data (only about 25% of the RNA-Seq detected transcripts were detected at the protein level). To further test the prediction we turned to *Drosophila* where an available high-resolution time-course allowed us to detect more LCI and HCI genes. Coupling the 3’ UTR isoform expression throughout embryogenesis (Sanfilippo et al. 2017) with total mRNA (Graveley et al. 2011) and protein expression data (Casas-Vila et al. 2017), we delineated LCI and HCI genes (see Methods) and studied the correlation between their transcription and protein levels in a *Drosophila melanogaster* embryonic time-course. As in *C. elegans*, we found that LCI genes exhibited reduced mRNA to protein expression correlation relative to HCI genes (Fig. 5B; *P*=0.00038, Mann-Whitney test; N=571, 189 for HCI and LCI, respectively). We further analyzed three extensive embryonic mRNA, protein and APA datasets from *Xenopus laevis* (Zhou et al. 2019; Peshkin et al. 2015) and observed similarly highly significant trends for this vertebrate species (Fig. 5C; *P*<10^−6^, Mann-Whitney test; N=1953, 996 for HCI and LCI, respectively). These results suggest that weak correlations between mRNA and protein may be in part explained by the existence of LCI genes with isoforms with distinct translation efficiencies.

**Fig. 5.**
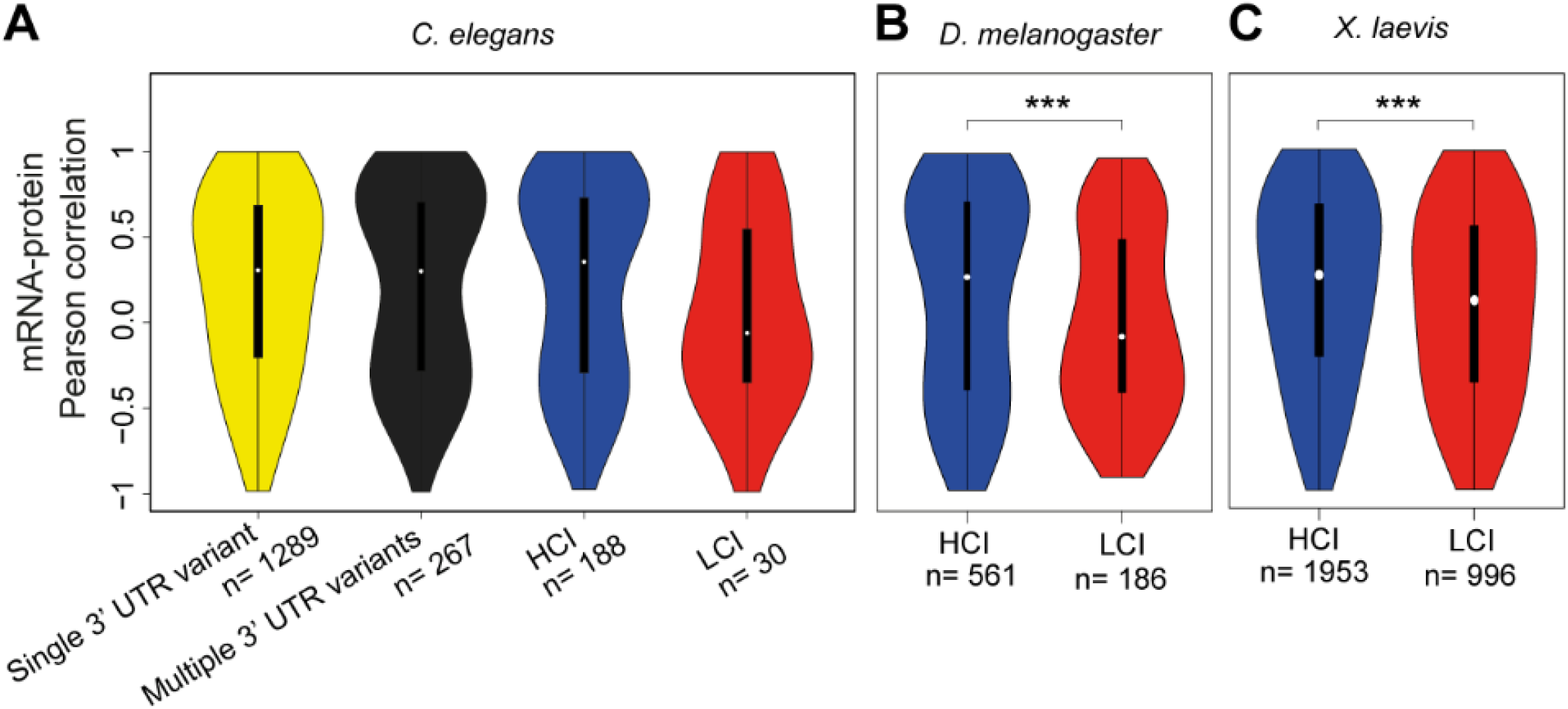
Genes with lowly correlated 3’ UTR isoforms (LCI genes) have a lower correspondence between total mRNA and protein expression. (A) Violin plots indicate the distribution of Pearson correlation coefficients between total mRNA and protein expression levels across developmental stages for *C. elegans* genes with one and multiple 3’ UTR isoforms, highly correlating (HCI), and lowly correlating isoforms (LCI). (B-C) Same as A, for LCI and HCI genes in *Drosophila melanogaster* (B) and *Xenopus laevis* (C) embryonic development.

## Discussion

The APA-seq approach allows for the extraction of an additional layer of data from CEL-Seq data. In addition to quantifying the expression of each gene, APA-seq reveals the alternative polyadenylation dynamics on a transcriptomic basis. APA-seq uses the CEL-Seq Read 2 to identify the gene of origin and then maps Read 1 to the respective gene sequence, instead of to the whole genome, enabling the use of even extremely low-quality sequencing data resulting from reading through protocol-conditioned poly-T stretches. Hence, by performing a subtle modification in data analysis without altering the actual CEL-Seq protocol, we were able to switch from analysis at the gene to the APA level. Although the presented data is based on CEL-Seq data, our method is in principle applicable to any RNA-Seq method that uses the polyA tail as anchor and performs paired end sequencing; including inDrop, 10X, and SMART-Seq (Klein et al. 2015; Picelli et al. 2013; Zheng et al. 2017). Other 3’ end sequencing methods have enabled important insights into the biological significance and mechanistic aspects of APA, though their experimental procedures require either relatively large amount of starting material or complex protocols combining several amplification steps (IVT and many PCR cycles) (Shepard et al. 2011; Derti et al. 2012; Yao et al. 2012; Lianoglou et al. 2013; Wilkening et al. 2013; Gruber et al. 2014; Nam et al. 2014; Li et al. 2015; Velten et al. 2015; Ye et al. 2018, Gupta et al. 2018).

In addition to APA-seq, two methods are currently available for assessing APA in low-input samples: BATSeq (Velten et al. 2015) and ScISOr-Seq (Gupta et al. 2018). BATSeq achieves single-cell APA isoform measurements. An important advantage of APA-seq relative to BATSeq is the formers high sensitivity, deriving from its use of CEL-Seq data; employing far less amplification and clean-up steps and PCR cycles, and a simple protocol and analysis. ScISOr-Seq also constitutes pioneering single-cell isoform work with the advantage of revealing the complete isoform due to its reliance on long-read sequencing. Relative to ScISOr-Seq, APA-Seq has the advantage of higher statistical confidence due to its reliance on deep Illumina sequencing. We highlight that APA-seq is not aimed towards the identification of polyA sites from scratch but rather we present it as an efficient method for quantifying the expression profiles of previously mapped isoforms. Furthermore, while we applied APA-seq here to single-embryo CEL-Seq data, it could in principle also be applied to single-cell data. As the protocol is relatively straight-forward omitting unnecessary clean-up and size-selection steps, the efficiency, complexity and accuracy is as high as that in CEL-Seq (Hashimshony et al. 2016, 2012). Applying APA-seq at single-cell resolution has many interesting applications, such as the study of population of cells undergoing cell fate specification, differentiation, and during tumorigenesis.

When studying the expression of alternative 3’ UTR isoforms throughout development, we found that the global transitions of isoform usage correspond to distinct developmental periods (Fig. 2). Our results are reminiscent of those of Lianoglou et al., who studied malignant transformation across human cell lines, and revealed a change in mRNA abundance levels of genes with a single 3’ UTR isoform, and, in genes with multiple 3’ UTR isoforms, a change in 3’ UTR isoform ratios (Lianoglou et al. 2013). A similar pattern emerged while comparing embryonic stem cells within differentiated tissues (Lianoglou et al. 2013). Our own results add an interesting layer to this field, by dissecting the temporal components of normal *C. elegans* embryonic development, providing insight into APA dynamics across distinct developmental stages. More generally, results are consistent with the accumulating evidence indicating that the APA regulatory mechanism is highly conserved across all eukaryotes, regardless of their morphological complexity (Ara et al. 2006; Wang et al. 2008; Shi 2012; Ulitsky et al. 2012; Velten et al. 2015; Hu et al. 2017).

3’ UTR isoforms are ubiquitous and considerable effort has been addressed towards understanding the distinct functional roles of different isoforms of the same gene (Tian and Manley 2017). Here, we report two general gene classes with 3’ UTR isoforms: those whose 3’ UTR isoforms correlate in their expression profiles across time and those that do not. Most genes (more than 70%) with alternatively polyadenylated isoforms exhibit a high correlation among their 3’ UTR isoforms. These highly correlated isoform genes (HCI) may be dominantly regulated at the level of overall transcription. In other words, the main factor influencing the distribution of the 3’ UTR isoforms usage is the intrinsic strength of their polyadenylation sites. Genes belonging to this class are often referred to as ‘dynamic genes’, which we previously showed to be enriched for developmental functions such as specification and differentiation (Levin et al. 2012; Levin et al. 2016). The HCI genes display a relative paucity of miRNA binding sites and higher concordance between total mRNA and protein levels (Figs. 4 and 5).

A rather small fraction of genes with multiple 3’ UTR isoforms (11%) show lowly correlated isoform expression across time. We have named these LCI genes and provide evidence that this gene class is under unique regulation. LCI genes show overall less dynamic total expression profiles; however, at the level of individual 3’ UTR isoforms they are highly dynamic (Fig. 3). LCI genes also exhibit several features which indicate a higher level of post-transcriptional regulation of 3’ UTR isoform usage. Post-transcriptional 3’ UTR-isoform regulation processes include miRNA mediated degradation and RNA-binding protein mediated stabilization or destabilization of mRNA molecules and control of translation efficiencies. 3’ UTRs of the LCI genes comprise significantly more miRNA binding sites and the 3’ UTR isoform ratio correlates well with the expression of a sub-group of miRNAs, many of which are known regulators of embryogenesis (Fig. 4). Consistently with these findings, the correlation between total mRNA and protein abundances is lower in LCI genes relative to HCI genes indicating that the 3’ UTR usage of this group of genes is tightly regulated.

Our results reveal principles of selective pressures on alternative polyadenylation. By studying APA dynamics over developmental time, we revealed two classes of genes with alternatively polyadenylated isoforms, the HCI and LCI. The LCI show the hallmarks of strong regulation on their 3’ UTR isoforms in the form of miRNA. Thus, we revealed here that a powerful litmus test for a functional distinction among 3’ UTR isoforms during a biological process (such as embryogenesis) is their discordant expression across time. As a corollary, genes with 3’ UTR isoforms showing correlated expression may represent biological noise of no functional consequence, as is common for other processes such as alternative splicing (Grishkevich and Yanai 2014). Collectively, our results characterize the regulatory principles of alternative polyadenylation and provide a context for the incorporation of specific posttranscriptional regulators such as miRNAs in the modeling of biological pathways.

## Methods

### Detection of polyadenylation site using CEL-Seq reads

We used our previously published *C. elegans* time-course data (GSE50548) sequenced using the CEL-Seq protocol and paired-end 100 bp sequencing mode. The CEL-Seq Read 2 insert was used to identify the gene by mapping reads to the reference genome (version WS230) using Bowtie 2 version 2.2.3 (Langmead and Salzberg 2012) with default parameters. The htseq-count algorithm (Anders et al. 2015) coupled with the genomic feature file was used to assign each individual Read 2 to its gene of origin. For each gene we extracted the whole gene coding sequence as well as the 5000 nucleotides downstream of the stop codon, or fewer than 5000 nucleotides if another gene was found in closer proximity. We truncated Read 1, removing the barcode sequence and the polyT stretch, leaving a sequence of approximately 70 nucleotides. We then used Bowtie 2 (Langmead and Salzberg 2012) in order to map Read 1 exclusively to the coding sequence of the identified gene. The maximum and minimum mismatch penalty (--mp MX,MN) parameters for Bowtie 2 were set to 2 and 1, respectively. To summarize the data for each sample, we counted all 3’-most mapping locations of truncated Read 1 to the respective genes up to a distance of 20 nucleotides upstream of the polyadenylation site. We predicted APA sites only for those genes passing a threshold of 20 mapped reads in at least two samples. For these genes we identified the peaks representing the polyadenylation sites by summarizing the last mapping coordinates of all the truncated Read 1 entities that mapped to a specific gene using the ‘findpeaks’ function in MATLAB. Peaks were required to be separated by at least 20 nucleotides in order to be considered as distinct. The height threshold for a peak was 5 reads, or 1/1000 of the total gene expression in a particular sample. We then filtered out possible spurious peaks that may have resulted from internal priming, by removing any peak whose downstream genomic sequence included any of the following nucleotide combinations: AAAA, AGAA, AAGA or AAAG (Gruber et al. 2016). The exact coordinate of any polyadenylation site was defined as the most 3’ coordinate of the respective peak. To validate the quality of our data, we compared our polyadenylation site annotation with a previously published *C. elegans* 3’ UTR annotation (Mangone et al. 2008). We performed this by measuring the difference between the mapping positions of polyadenylation sites from both data sets (see Fig. 1D).

### 3’ UTR isoform expression throughout *C. elegans* development

Expression data for each sample was obtained by counting all Read 1 sequences whose 3’-most mapping location mapped up to a distance of 20 nucleotides upstream of the polyadenylation site. The raw expression data was then converted to transcripts per million (tpm) by dividing by the total reads and multiplying by one million. We worked with log_10_ values unless otherwise noted. The profiles were further smoothed by computing a running average over 5 time points. To study dynamics of 3’ UTR isoform usage throughout the *C. elegans* embryonic time-course, we first filtered genes on overall expression, keeping only those in the upper 85 percentile of the sum of expression throughout the time-course. Ratios between the two most highly expressed 3’ UTR isoforms where calculated for all genes and all stages by dividing the expression levels (tpm) of the short by the levels of the long 3’ UTR isoform. Only genes whose 3’ UTR isoform ratios across time differed by at least a factor of three were kept for further analysis ending up with 305 genes. Significance of the changes in the ratios during the transition between successive periods were calculated using Student’s t-test on all ratios of one and the ratios of the successive period. *P*-values and fold changes are shown in Fig. 2B. To generate temporally sorted expression profiles, we used “ZAVIT” as previously described (Levin et al. 2016; Zalts and Yanai 2017).

### Categorization of genes by APA behavior

To determine whether total gene expression is considered static or dynamic throughout development, we used the ratio of minimum to maximum and as validation the interquartile range (IQR) of expression levels across time. To quantify 3’ UTR variant expression deviations, we calculated Pearson correlation for the expression pattern of the two, or more, 3’ UTR isoforms. For genes with more than two isoforms (<1% of expressed genes), we used the minimal Pearson correlation between any pair of isoforms. Highly correlated 3’ UTR isoforms (HCI) are those with *r*>0.7, while lowly correlated 3’ UTR isoforms (LCI) are those with *r*
<0.3.

### miRNA target analysis

3’ UTR sequences were identified according to APA-seq. After removing gene coding sequences using WormBase annotation, we annotated miRNA binding sites by searching for the basic seed match sequence within the 3’ UTR, i.e. the complementary sequence for nucleotides 2-7 of the miRNA (Ambros 2004; Bartel 2004). We disregarded other parameters such as type of seed match or complementarity outside of the seed region. We then counted the number of miRNA binding sites for each gene subgroup (LCI and HCI genes), and used Student’s t-test to determine the significance of the difference in the number of miRNA binding sites between the LCI and HCI groups. To determine the effect specific miRNAs have on their targets, in genes with multiple 3’ UTR isoforms, we calculated Pearson correlation between the isoform expression ratio of the two most highly expressed isoforms, and the miRNA expression profile (Avital et al. 2017). For this analysis we examined only dynamically expressed miRNAs, whose expression is above 250 transcripts overall across the timepoints. To compare the miRNA dataset with our APA-seq dataset, we examined only matching timepoints to our time-course and used the Matlab ‘imresize’ function followed by smoothing to stretch the miRNA expression data.

### mRNA-protein correlation analysis

RNA-Seq and Silac data of different developmental stages in *C. elegans* was downloaded from Grün et al. (Grün et al. 2014). Replicates were averaged and data was normalized by division by the sum of all genes for the specific samples. Correlation between rpkm (mRNA) and Silac (protein) expression values using Pearson correlation was computed only for genes with rpkm and silac values for at least three stages. A similar approach was used to calculate the correlation between RNA and protein levels for the *Drosophila melanogaster* time-course. APA data of an embryonic time-course was downloaded from Sanfilippo et al (Sanfilippo et al. 2017); 3’ UTR isoform ratio was analyzed similarly to our time-course data yielding correlation between 3’ UTR isoforms for all multi-isoform genes. Highly correlated 3’ UTR isoforms (HCI) are those with *r*>0.7, while lowly correlated 3’ UTR isoforms (LCI) are those with *r*<0.3, coherent with the thresholds used for our dataset. Total mRNA and protein expression levels were downloaded from Graveley et al. (Graveley et al. 2011) and Casas-Vila et al. (Casas-Vila et al. 2017), respectively. The two datasets were integrated by correlating mRNA (rpkm) and protein (lfq) expression levels using Pearson correlation. The same procedure was used to analyze the data for *Xenopus laevis*. mRNA RNA-Seq and protein LFQ data for embryonic time points were downloaded from (Peshkin et al. 2015) and APA data from (Zhou et al. 2019). For the datasets from all three species, the Mann-Whitney test was used to determine if LCI genes show different mRNA-protein expression correlations from HCI genes.

## Acknowledgements

We thank Eitan Winter and Martin Feder for their assistance throughout this project. We also acknowledge assistance from the Technion Genome Center.

## Author contributions

N.M., T.H., and I.Y. conceived and designed the project. N.M. led the development of the APA-seq approach. M.L., H.Z., N.M., and I.Y. analyzed the 3’ UTR isoform data. H.Z. contributed the miRNA analysis and comparison with previous annotations. M.L. contributed the developmental APA dynamics, mRNA-protein correlation and RNA-binding-protein analyses. M.L., H.Z., N.M. and I.Y. led the interpretation of the data. M.L., H.Z., N.M., and I.Y. drafted the manuscript. M.L. is supported by a Humboldt-Bayer research fellowship.

## Disclosure declaration

The authors declare that they have no conflicts of interest.

## References

Alt FW, Bothwell AL, Knapp M, Siden E, Mather E, Koshland M, Baltimore D. 1980. Synthesis of secreted and membrane-bound immunoglobulin mu heavy chains is directed by mRNAs that differ at their 3’ ends. Cell 20: 293–301.

Ambros V. 2004. The functions of animal microRNAs. Nature 431: 350–355.

Anders S, Pyl PT, Huber W. 2015. HTSeq--a Python framework to work with high-throughput sequencing data. Bioinformatics 31: 166–169.

Ara T, Lopez F, Ritchie W, Benech P, Gautheret D. 2006. Conservation of alternative polyadenylation patterns in mammalian genes. BMC Genomics 7: 189.

Avital G, Starvaggi França G, Yanai I. 2017. Bimodal evolutionary developmental miRNA program in animal embryogenesis. Mol Biol Evol doi:10.1093/molbev/msx316.

Barrett LW, Fletcher S, Wilton SD. 2012. Regulation of eukaryotic gene expression by the untranslated gene regions and other non-coding elements. Cell Mol Life Sci 69: 3613–3634.

Bartel DP. 2004. MicroRNAs: genomics, biogenesis, mechanism, and function. Cell 116: 281–297.

Berkovits BD, Mayr C. 2015. Alternative 3’ UTRs act as scaffolds to regulate membrane protein localization. Nature 522: 363–367.

Blazie SM, Babb C, Wilky H, Rawls A, Park JG, Mangone M. 2015. Comparative RNA-Seq analysis reveals pervasive tissue-specific alternative polyadenylation in Caenorhabditis elegans intestine and muscles. BMC Biol 13: 4.

Blazie SM, Geissel HC, Wilky H, Joshi R, Newbern J, Mangone M. 2017. Alternative Polyadenylation Directs Tissue-Specific miRNA Targeting in Caenorhabditis elegans Somatic Tissues. Genetics 206: 757–774.

Casas-Vila N, Bluhm A, Sayols S, Dinges N, Dejung M, Altenhein T, Kappei D, Altenhein B, Roignant J-Y, Butter F. 2017. The developmental proteome of Drosophila melanogaster. Genome Res 27: 1273–1285.

Chen W, Jia Q, Song Y, Fu H, Wei G, Ni T. 2017. Alternative Polyadenylation: Methods, Findings, and Impacts. Genomics Proteomics Bioinformatics 15: 287–300.

Consortium CeS. 1998. Genome sequence of the nematode C. elegans: a platform for investigating biology. Science 282: 2012–2018.

de Lucas S, Oliveros JC, Chagoyen M, Ortín J. 2014. Functional signature for the recognition of specific target mRNAs by human Staufen1 protein. Nucleic Acids Res 42: 4516–4526.

Derti A, Garrett-Engele P, Macisaac KD, Stevens RC, Sriram S, Chen R, Rohl CA, Johnson JM, Babak T. 2012. A quantitative atlas of polyadenylation in five mammals. Genome Res 22: 1173–1183.

Early P, Rogers J, Davis M, Calame K, Bond M, Wall R, Hood L. 1980. Two mRNAs can be produced from a single immunoglobulin mu gene by alternative RNA processing pathways. Cell 20: 313–319.

Elkon R, Ugalde AP, Agami R. 2013. Alternative cleavage and polyadenylation: extent, regulation and function. Nat Rev Genet 14: 496–506.

Goldstrohm AC, Hall TMT, McKenney KM. 2018. Post-transcriptional Regulatory Functions of Mammalian Pumilio Proteins. Trends Genet 34: 972–990.

Graveley BR, Brooks AN, Carlson JW, Duff MO, Landolin JM, Yang L, Artieri CG, van Baren MJ, Boley N, Booth BW et al. 2011. The developmental transcriptome of Drosophila melanogaster. Nature 471: 473–479.

Grishkevich V, Yanai I. 2014. Gene length and expression level shape genomic novelties. Genome Res 24: 1497–1503.

Gruber AJ, Schmidt R, Gruber AR, Martin G, Ghosh S, Belmadani M, Keller W, Zavolan M. 2016. A comprehensive analysis of 3’ end sequencing data sets reveals novel polyadenylation signals and the repressive role of heterogeneous ribonucleoprotein C on cleavage and polyadenylation. Genome Res 26: 1145–1159.

Gruber AR, Martin G, Müller P, Schmidt A, Gruber AJ, Gumienny R, Mittal N, Jayachandran R, Pieters J, Keller W et al. 2014. Global 3’ UTR shortening has a limited effect on protein abundance in proliferating T cells. Nat Commun 5: 5465.

Grün D, Kirchner M, Thierfelder N, Stoeckius M, Selbach M, Rajewsky N. 2014. Conservation of mRNA and protein expression during development of C. elegans. Cell Rep 6: 565–577.

Hashimshony T, Senderovich N, Avital G, Klochendler A, de Leeuw Y, Anavy L, Gennert D, Li S, Livak KJ, Rozenblatt-Rosen O et al. 2016. CEL-Seq2: sensitive highly-multiplexed single-cell RNA-Seq. Genome Biol 17: 77.

Hashimshony T, Wagner F, Sher N, Yanai I. 2012. CEL-Seq: single-cell RNA-Seq by multiplexed linear amplification. Cell Rep 2: 666–673.

Hillier LW, Reinke V, Green P, Hirst M, Marra MA, Waterston RH. 2009. Massively parallel sequencing of the polyadenylated transcriptome of C. elegans. Genome Res 19: 657–666.

Hu W, Li S, Park JY, Boppana S, Ni T, Li M, Zhu J, Tian B, Xie Z, Xiang M. 2017. Dynamic landscape of alternative polyadenylation during retinal development. Cellular and molecular life sciences: CMLS 74: 1721–1739.

Jan CH, Friedman RC, Ruby JG, Bartel DP. 2011. Formation, regulation and evolution of Caenorhabditis elegans 3’UTRs. Nature 469: 97–101.

Ji Z, Lee JY, Pan Z, Jiang B, Tian B. 2009. Progressive lengthening of 3’ untranslated regions of mRNAs by alternative polyadenylation during mouse embryonic development. Proc Natl Acad Sci USA 106: 7028–7033.

Kamath RS, Fraser AG, Dong Y, Poulin G, Durbin R, Gotta M, Kanapin A, Le Bot N, Moreno S, Sohrmann M et al. 2003. Systematic functional analysis of the Caenorhabditis elegans genome using RNAi. Nature 421: 231–237.

Keene JD. 2007. RNA regulons: coordination of post-transcriptional events. Nat Rev Genet 8: 533–543.

Khraiwesh B, Salehi-Ashtiani K. 2017. Alternative Poly(A) Tails Meet miRNA Targeting in Caenorhabditis elegans. Genetics 206: 755–756.

Kim YK, Furic L, Desgroseillers L, Maquat LE. 2005. Mammalian Staufen1 recruits Upf1 to specific mRNA 3’UTRs so as to elicit mRNA decay. Cell 120: 195–208.

Kimble J, Hirsh D. 1979. The postembryonic cell lineages of the hermaphrodite and male gonads in Caenorhabditis elegans. Dev Biol 70: 396–417.

Klein AM, Mazutis L, Akartuna I, Tallapragada N, Veres A, Li V, Peshkin L, Weitz DA, Kirschner MW. 2015. Droplet Barcoding for Single-Cell Transcriptomics Applied to Embryonic Stem Cells. Cell 161: 1187–1201. http://www.ncbi.nlm.nih.gov/pubmed/26000487 (Accessed May 21, 2015).

Lamm AT, Stadler MR, Zhang H, Gent JI, Fire AZ. 2011. Multimodal RNA-seq using single-strand, double-strand, and CircLigase-based capture yields a refined and extended description of the C. elegans transcriptome. Genome Res 21: 265–275.

Langmead B, Salzberg SL. 2012. Fast gapped-read alignment with Bowtie 2. Nat Methods 9: 357–359.

Lee C-YS, Lu T, Seydoux G. 2017. Nanos promotes epigenetic reprograming of the germline by down-regulation of the THAP transcription factor LIN-15B. Elife 6.

Levin M, Anavy L, Cole AG, Winter E, Mostov N, Khair S, Senderovich N, Kovalev E, Silver DH, Feder M et al. 2016. The mid-developmental transition and the evolution of animal body plans. Nature 531: 637–641.

Levin M, Hashimshony T, Wagner F, Yanai I. 2012. Developmental milestones punctuate gene expression in the Caenorhabditis embryo. Dev Cell 22: 1101–1108.

Li W, You B, Hoque M, Zheng D, Luo W, Ji Z, Park JY, Gunderson SI, Kalsotra A, Manley JL et al. 2015. Systematic profiling of poly(A)+ transcripts modulated by core 3’ end processing and splicing factors reveals regulatory rules of alternative cleavage and polyadenylation. PLoS Genet 11: e1005166.

Li Y, Sun Y, Fu Y, Li M, Huang G, Zhang C, Liang J, Huang S, Shen G, Yuan S et al. 2012. Dynamic landscape of tandem 3’ UTRs during zebrafish development. Genome Res 22: 1899–1906.

Lianoglou S, Garg V, Yang JL, Leslie CS, Mayr C. 2013. Ubiquitously transcribed genes use alternative polyadenylation to achieve tissue-specific expression. Genes Dev 27: 2380–2396.

Lutz CS, Moreira A. 2011. Alternative mRNA polyadenylation in eukaryotes: an effective regulator of gene expression. Wiley Interdiscip Rev RNA 2: 22–31.

Mangone M, Macmenamin P, Zegar C, Piano F, Gunsalus KC. 2008. UTRome.org: a platform for 3’UTR biology in C. elegans. Nucleic Acids Res 36: D57–62.

Mangone M, Manoharan AP, Thierry-Mieg D, Thierry-Mieg J, Han T, Mackowiak SD, Mis E, Zegar C, Gutwein MR, Khivansara V et al. 2010. The landscape of C. elegans 3’UTRs. Science 329: 432–435.

Marco A, Hooks K, Griffiths-Jones S. 2012. Evolution and function of the extended miR-2 microRNA family. RNA Biol 9: 242–248.

Mazumder B, Seshadri V, Fox PL. 2003. Translational control by the 3’-UTR: the ends specify the means. Trends Biochem Sci 28: 91–98.

McKay SJ, Johnsen R, Khattra J, Asano J, Baillie DL, Chan S, Dube N, Fang L, Goszczynski B, Ha E et al. 2003. Gene expression profiling of cells, tissues, and developmental stages of the nematode C. elegans. Cold Spring Harb Symp Quant Biol 68: 159–169.

Micklem DR, Adams J, Grünert S, St Johnston D. 2000. Distinct roles of two conserved Staufen domains in oskar mRNA localization and translation. EMBO J 19: 1366–1377.

Mitchell SF, Parker R. 2014. Principles and properties of eukaryotic mRNPs. Mol Cell 54: 547–558.

Müller-McNicoll M, Neugebauer KM. 2013. How cells get the message: dynamic assembly and function of mRNA-protein complexes. Nat Rev Genet 14: 275–287.

Nam J-W, Rissland OS, Koppstein D, Abreu-Goodger C, Jan CH, Agarwal V, Yildirim MA, Rodriguez A, Bartel DP. 2014. Global analyses of the effect of different cellular contexts on microRNA targeting. Mol Cell 53: 1031–1043.

Newman-Smith ED, Rothman JH. 1998. The maternal-to-zygotic transition in embryonic patterning of Caenorhabditis elegans. Curr Opin Genet Dev 8: 472–480.

Ozsolak F, Kapranov P, Foissac S, Kim SW, Fishilevich E, Monaghan AP, John B, Milos PM. 2010. Comprehensive polyadenylation site maps in yeast and human reveal pervasive alternative polyadenylation. Cell 143: 1018–1029.

Pagano JM, Farley BM, Essien KI, Ryder SP. 2009. RNA recognition by the embryonic cell fate determinant and germline totipotency factor MEX-3. Proc Natl Acad Sci USA 106: 20252–20257.

Peshkin L, Wühr M, Pearl E, Haas W, Freeman RM, Gerhart JC, Klein AM, Horb M, Gygi SP, Kirschner MW. 2015. On the Relationship of Protein and mRNA Dynamics in Vertebrate Embryonic Development. Dev Cell 35: 383–94. https://linkinghub.elsevier.com/retrieve/pii/S1534580715006577 (Accessed April 6, 2019).

Peterson SM, Thompson JA, Ufkin ML, Sathyanarayana P, Liaw L, Congdon CB. 2014. Common features of microRNA target prediction tools. Front Genet 5: 23.

Picelli S, Björklund ÅK, Faridani OR, Sagasser S, Winberg G, Sandberg R. 2013. Smart-seq2 for sensitive full-length transcriptome profiling in single cells. Nat Methods 10: 1096–8. http://www.ncbi.nlm.nih.gov/pubmed/24056875 (Accessed July 12, 2014)

Pichon X, Wilson LA, Stoneley M, Bastide A, King HA, Somers J, Willis AEE. 2012. RNA binding protein/RNA element interactions and the control of translation. Curr Protein Pept Sci 13: 294–304.

Prasad A, Porter DF, Kroll-Conner PL, Mohanty I, Ryan AR, Crittenden SL, Wickens M, Kimble J. 2016. The PUF binding landscape in metazoan germ cells. RNA 22: 1026–1043.

Ramani AK, Nelson AC, Kapranov P, Bell I, Gingeras TR, Fraser AG. 2009. High resolution transcriptome maps for wild-type and nonsense-mediated decay-defective Caenorhabditis elegans. Genome Biol 10: R101.

Rogers J, Early P, Carter C, Calame K, Bond M, Hood L, Wall R. 1980. Two mRNAs with different 3’ ends encode membrane-bound and secreted forms of immunoglobulin mu chain. Cell 20: 303–312.

Sanfilippo P, Wen J, Lai EC. 2017. Landscape and evolution of tissue-specific alternative polyadenylation across Drosophila species. Genome Biol 18: 229.

Setzer DR, McGrogan M, Nunberg JH, Schimke RT. 1980. Size heterogeneity in the 3’ end of dihydrofolate reductase messenger RNAs in mouse cells. Cell 22: 361–370.

Shepard PJ, Choi E-A, Lu J, Flanagan LA, Hertel KJ, Shi Y. 2011. Complex and dynamic landscape of RNA polyadenylation revealed by PAS-Seq. RNA 17: 761–772.

Shi Y. 2012. Alternative polyadenylation: new insights from global analyses. RNA 18: 2105–2117.

Shin H, Hirst M, Bainbridge MN, Magrini V, Mardis E, Moerman DG, Marra MA, Baillie DL, Jones SJM. 2008. Transcriptome analysis for Caenorhabditis elegans based on novel expressed sequence tags. BMC Biol 6: 30.

Smibert P, Miura P, Westholm JO, Shenker S, May G, Duff MO, Zhang D, Eads BD, Carlson J, Brown JB et al. 2012. Global patterns of tissue-specific alternative polyadenylation in Drosophila. Cell Rep 1: 277–289.

Smith CJ, Watson JD, Spencer WC, O’Brien T, Cha B, Albeg A, Treinin M, Miller DM. 2010. Time-lapse imaging and cell-specific expression profiling reveal dynamic branching and molecular determinants of a multi-dendritic nociceptor in C. elegans. Dev Biol 345: 18–33.

Sulston JE, Horvitz HR. 1977. Post-embryonic cell lineages of the nematode, Caenorhabditis elegans. Dev Biol 56: 110–156.

Szostak E, Gebauer F. 2013. Translational control by 3’-UTR-binding proteins. Brief Funct Genomics 12: 58–65.

Tadros W, Goldman AL, Babak T, Menzies F, Vardy L, Orr-Weaver T, Hughes TR, Westwood JT, Smibert CA, Lipshitz HD. 2007a. SMAUG is a major regulator of maternal mRNA destabilization in Drosophila and its translation is activated by the PAN GU kinase. Dev Cell 12: 143–155.

Tadros W, Lipshitz HD. 2009. The maternal-to-zygotic transition: a play in two acts. Development 136: 3033–3042.

Tadros W, Westwood JT, Lipshitz HD. 2007b. The mother-to-child transition. Dev Cell 12: 847–849.

Tian B, Hu J, Zhang H, Lutz CS. 2005. A large-scale analysis of mRNA polyadenylation of human and mouse genes. Nucleic Acids Res 33: 201–212.

Tian B, Manley JL. 2017. Alternative polyadenylation of mRNA precursors. Nat Rev Mol Cell Biol 18: 18–30.

Ulitsky I, Shkumatava A, Jan CH, Subtelny AO, Koppstein D, Bell GW, Sive H, Bartel DP. 2012. Extensive alternative polyadenylation during zebrafish development. Genome Res 22: 2054–2066.

Velten L, Anders S, Pekowska A, Järvelin AI, Huber W, Pelechano V, Steinmetz LM. 2015. Single-cell polyadenylation site mapping reveals 3’ isoform choice variability. Mol Syst Biol 11: 812.

Walser CB, Lipshitz HD. 2011. Transcript clearance during the maternal-to-zygotic transition. Curr Opin Genet Dev 21: 431–443.

Wang ET, Sandberg R, Luo S, Khrebtukova I, Zhang L, Mayr C, Kingsmore SF, Schroth GP, Burge CB. 2008. Alternative isoform regulation in human tissue transcriptomes. Nature 456: 470–476.

Wilkening S, Pelechano V, Järvelin AI, Tekkedil MM, Anders S, Benes V, Steinmetz LM. 2013. An efficient method for genome-wide polyadenylation site mapping and RNA quantification. Nucleic Acids Res 41: e65.

Wormington M. 1994. Unmasking the role of the 3’ UTR in the cytoplasmic polyadenylation and translational regulation of maternal mRNAs. Bioessays 16: 533–535.

Yao C, Biesinger J, Wan J, Weng L, Xing Y, Xie X, Shi Y. 2012. Transcriptome-wide analyses of CstF64-RNA interactions in global regulation of mRNA alternative polyadenylation. Proc Natl Acad Sci USA 109: 18773–18778.

Ye C, Long Y, Ji G, Li QQ, Wu X. 2018. APAtrap: identification and quantification of alternative polyadenylation sites from RNA-seq data. Bioinformatics 34: 1841–1849.

Zalts H, Yanai I. 2017. Developmental constraints shape the evolution of the nematode mid-developmental transition. Nature Ecology & Evolution 1: s41559-41017-40113-41017.

Zhao W, Pollack JL, Blagev DP, Zaitlen N, McManus MT, Erle DJ. 2014. Massively parallel functional annotation of 3’ untranslated regions. Nat Biotechnol 32: 387–391.

Zheng GXY, Terry JM, Belgrader P, Ryvkin P, Bent ZW, Wilson R, Ziraldo SB, Wheeler TD, McDermott GP, Zhu J, et al. 2017. Massively parallel digital transcriptional profiling of single cells. Nat Commun 8: 14049.

Zhou X, Zhang Y, Michal JJ, Qu L, Zhang S, Wildung MR, Du W, Pouchnik DJ, Zhao H, Xia Y, et al. 2019. Alternative polyadenylation coordinates embryonic development, sexual dimorphism and longitudinal growth in Xenopus tropicalis. Cell Mol Life Sci. http://link.springer.com/10.1007/s00018-019-03036-1 (Accessed April 6, 2019).

