## Supplementary Figures for "Developmental dynamics are a proxy for selective pressures on alternatively polyadenylated isoforms"

### Contents

**Supplemental Figures S1-S5**

**Supplemental Tables S1-S2**

### Supplemental Figures

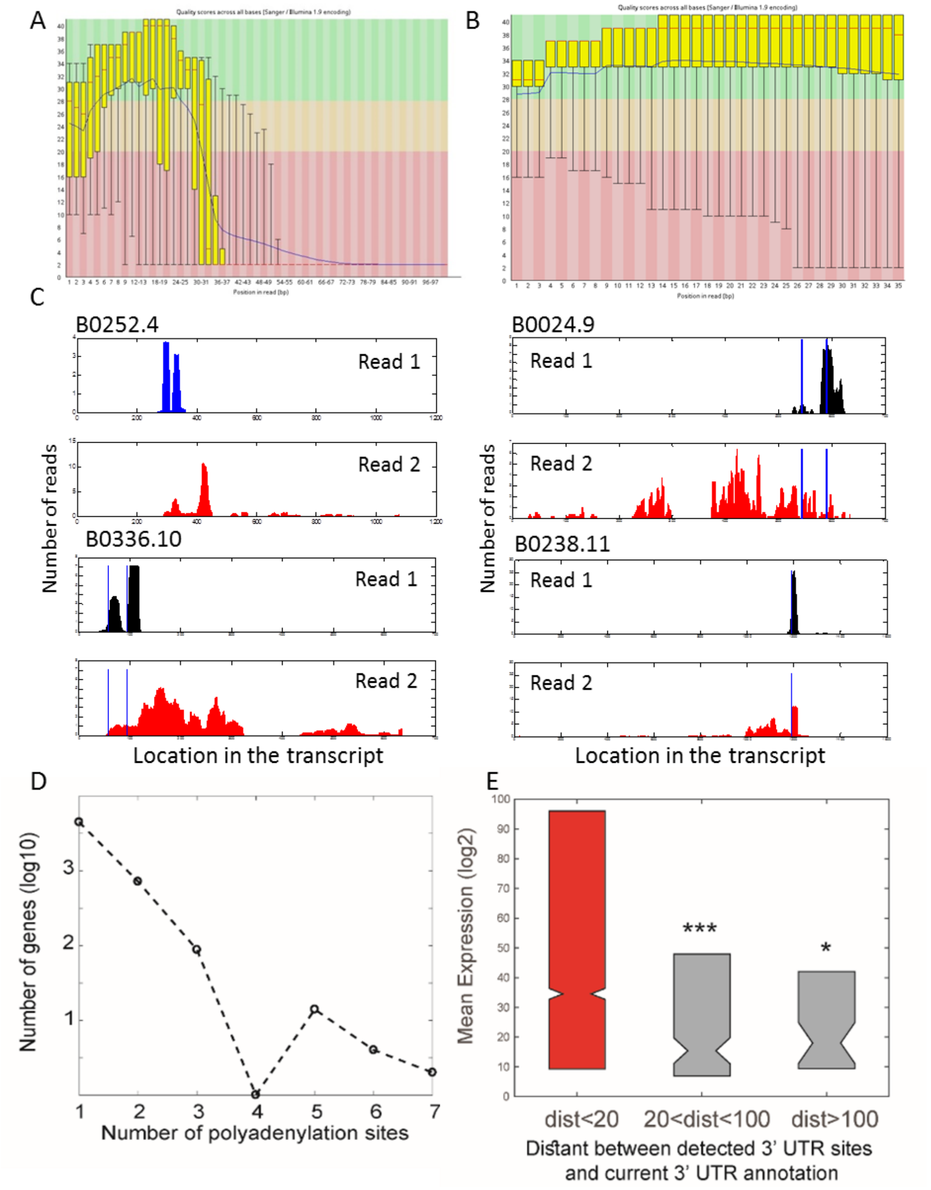


**Supplemental Figure S1. APA-seq measures expression levels of distinct 3' UTR isoforms.** (A) Read 1 quality scores across all bases. Note the fall in quality due to the sequencing of a polyT stretch. (B) Read 2 quality scores across all bases. (C) For the indicated genes, the distributions of Read 1 and Read 2 mappings are shown. Blue vertical lines indicate the location of known polyadenylation sites. (D) Histogram of the number of genes with different numbers of alternative 3' UTR isoforms. (E) Genes with 3' UTR isoforms that do not fully correspond to the commonly used 3' UTR isoform annotation *elegans* (Mangone et al. 2008) (5% of alternative isoforms found by APA-seq method) show significantly lower mean expression levels than the high confidence set.

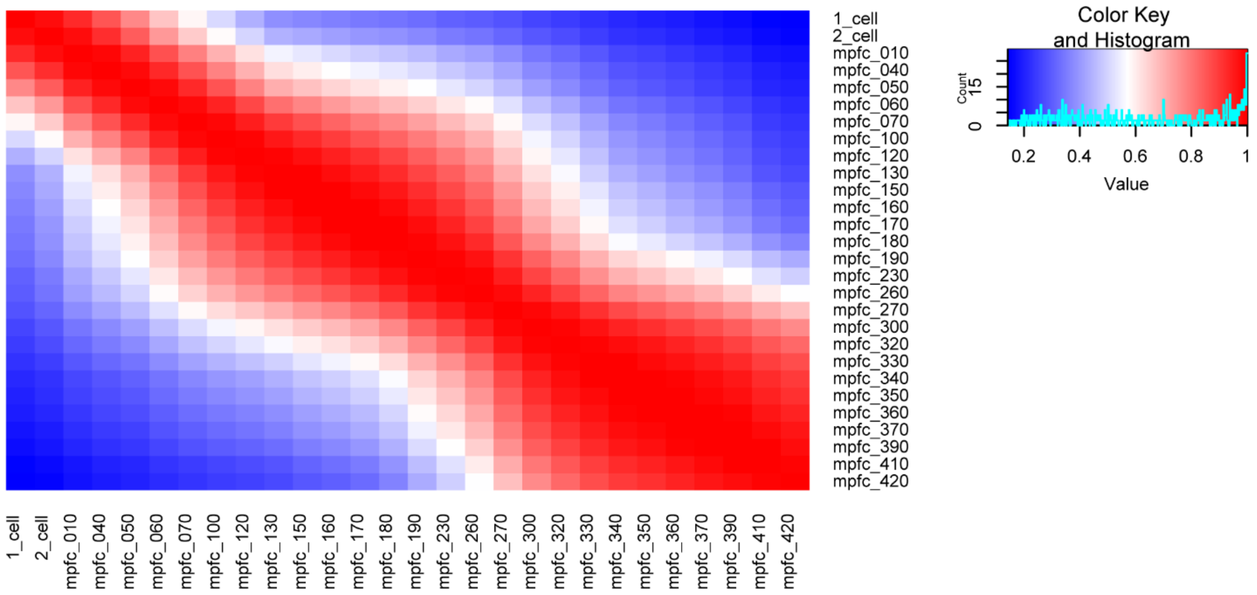


**Supplemental Figure S2. Heatmap of Pearson correlation coefficients of all pairwise stage 3' UTR isoform expression (tpm) comparisons.** 3' UTR isoform expression correlations show high similarities between successive stages. mpfc=minutes past 4-cell stage.

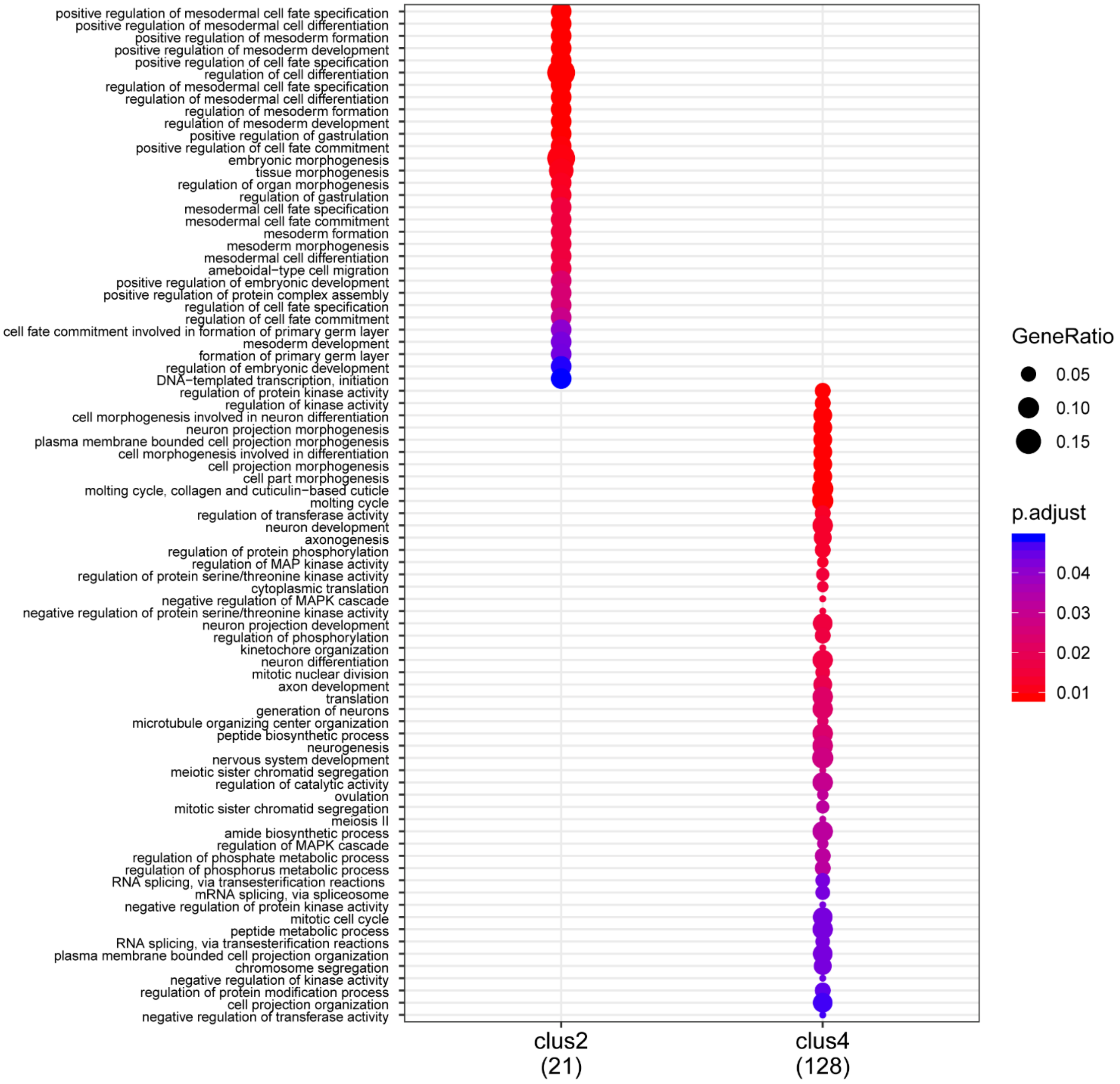


**Supplemental Figure S3. Gene Ontology (GO) enrichment analysis of 3' UTR isoform ratio clusters (see Fig. 2D).** GO terms of the 'biological process' category enriched in clusters determined in Fig. 2D. The size of the dots corresponds to the ratio of genes included in each term out of total number of genes in cluster. Color corresponds to adjusted *P*-value (hypergeometric distribution, Benjamini-Hochberg adjustment). Cluster 1 and 3 did not show any enrichment.

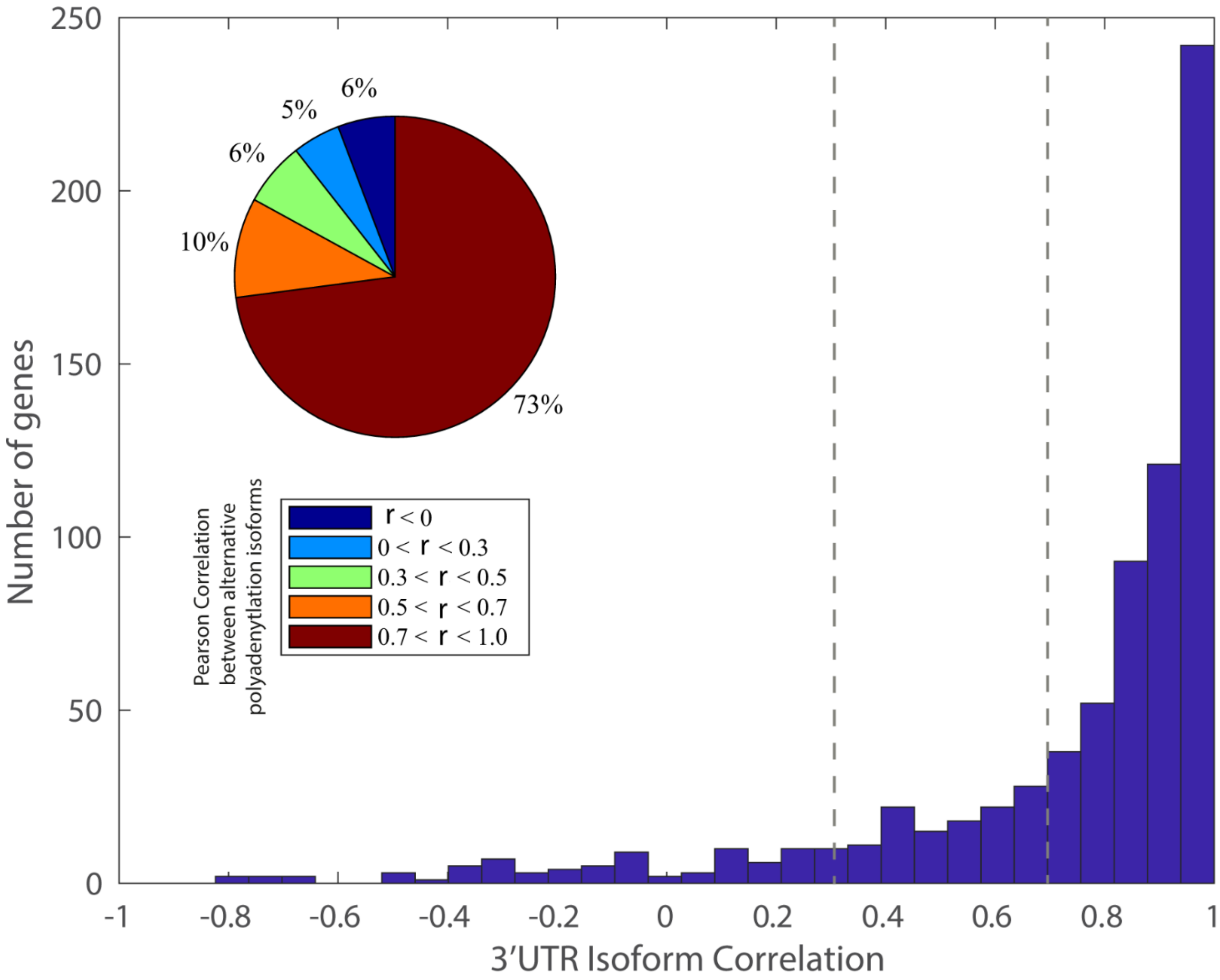


**Supplemental Figure S4. Histogram and pie chart indicating the distribution of 3' UTR isoform expression correlations.**

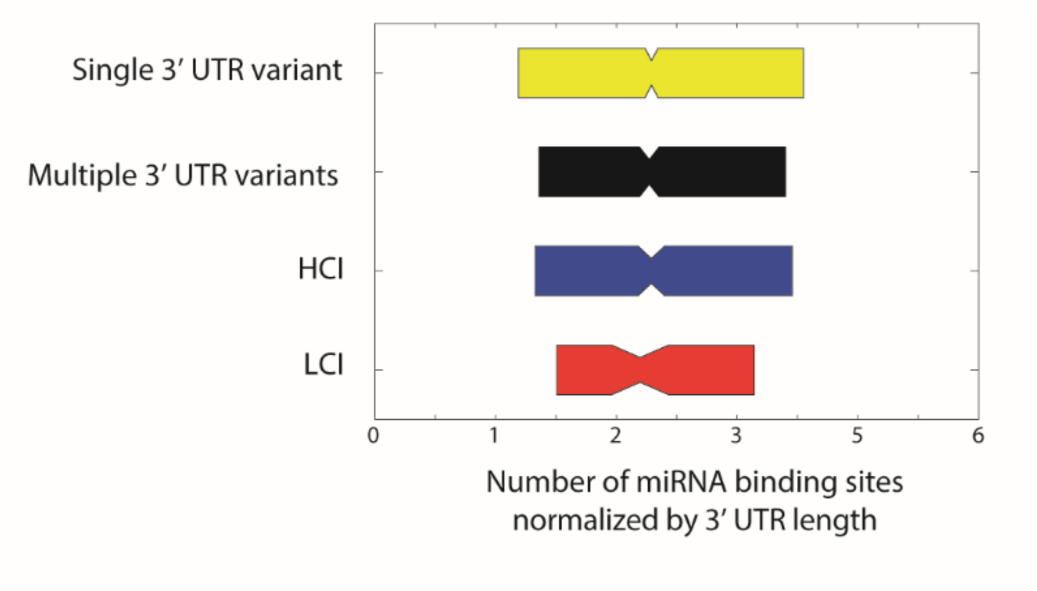


**Supplemental Figure S5. Concentration of miRNA binding sites across gene groups.** The boxplots indicate the number of miRNA binding sites normalized by 3' UTR length in genes with single or multiple 3' UTR isoforms and highly correlating isoforms (HCI) or lowly correlating isoforms (LCI) in yellow, black, blue and red, respectively. No significant difference was detected between genes of the different groups, when normalizing the amount of miRNA binding sites to 3' UTR length. Normalizing the number of miRNA seed matches to UTR length indicates that genes with multiple, lowly correlated 3' UTR isoforms have longer 3' UTRs but not a higher density of miRNA binding sites.

### Supplemental Tables

**Supplemental Table S1.** Expression levels (tpm) and annotation details retrieved by APA-seq.

**Supplemental Table S2.** Dynamically expressed miRNA correlation statistics.
